# High spontaneous integration rates of end-modified linear DNAs upon mammalian cell transfection

**DOI:** 10.1101/2022.12.19.521118

**Authors:** Samuel Lim, R. Rogers Yocum, Pamela A Silver, Jeffrey C Way

## Abstract

In gene therapy, potential integration of therapeutic transgene into host cell genomes is a serious risk that can lead to insertional mutagenesis and tumorigenesis. Viral vectors are often used as the gene delivery vehicle, but they are prone to undergoing integration events. More recently, non-viral delivery of linear DNAs having modified geometry such as closed-end linear duplex DNA (CELiD) have shown promise as an alternative, due to prolonged transgene expression and less cytotoxicity. However, whether such modified-end linear DNAs can also provide a safe, non-integrating gene transfer remains unanswered. Herein, we provide a systematic comparison of genomic integration frequency upon transfection of cells with expression vectors in the forms of circular plasmid, unmodified linear DNA, CELiD, and Streptavidin-conjugated blocked-end linear DNA. All of these forms of linear DNA resulted in a high fraction of the cells being stably transfected – between 10% and 20% of the initially transfected cells, with CELiDs showing the highest rates of integration. These results indicate that blocking the ends of linear DNA is insufficient to prevent integration. Moreover, our analysis suggest that conventional AAV-based gene therapy may be highly susceptible to integration, which is consistent with recent findings from long-term clinical studies.

## Introduction

Gene therapy aims to treat patients with genetic disease by introducing genetic material into cells to repair defective cellular functions. [1, 2] Traditionally, viral vectors such as retrovirus or lentivirus have been preferred as the gene delivery vehicles due to their high transfection efficiency. [3] However, viral vector-based methods suffer from a critical safety concern, regarding potential integration of DNA into a host cell chromosome, which may result in activation of oncogenes or knockout of tumor suppressor genes. [4] Indeed, there were multiple reports of patients developing oncogenesis during the early trials of gene therapy using viral vector delivery. [5, 6] While adeno-associated virus (AAV) vectors display less propensity to integrate, several studies indicate that the risk may still exist. [7-9] Because of such potential danger, the current scope of gene therapy is limited primarily to the treatment of cancer or rare genetic disorders, where patients and their caregivers are more likely to accept the risks.

Reflecting recent efforts to address the limits of viral vectors, there is a growing interest in non-viral delivery of nucleic acids. [10] Despite its simplicity and ease of production, circular plasmid DNAs may allow chromosomal integration through crossover events. Linear DNAs, including those derived from cleavage of circular DNAs after entering a cell, undergo non-homologous end joining (NHEJ) to be randomly integrated into the host cell genome, while being prone to exonuclease activity *in vivo*. Recently, close-ended linear duplex DNA (CELiD) that consists of double-stranded DNA molecules with covalently closed terminal hairpins have been reported. [11] A type of CELiD is generated as an intermediate of AAV replication in eukaryotic cells, and such structures are in effect a double stranded AAV genome capped with hairpin-forming palindromic terminal regions. [12] When CELiDs and circular plasmids were injected into a mouse, both forms of DNA were trafficked into the liver, but the CELiDs promoted extended and more stable expression of the transgenes. [11] This observation highlights the promising potential of end-modified linear DNA as an alternative non-viral vector for gene therapy. However, the question of whether this strategy can provide a safe, non-integrating gene delivery remains uncertain. While modifying DNA ends is known to hinder random end-joining events, [13-15] extended transgenic expression by CELiDs may result from a higher tendency for integration.

Herein, we report a comparative study on the genomic integration of linear DNAs having different types of modified ends, and investigate whether blocking DNA ends improves the safety of gene delivery. Specifically, we synthesized a novel blocked-end linear DNA using streptavidin-biotin bioconjugation chemistry [16], and subsequently compared its propensity for integration to those of CELiD, unmodified linear and circular DNAs. Upon transfection, we cultured the cells in the absence of a drug selection over extended time, during which transiently transfected genes would get diluted away and any remaining stable gene expression could be attributed to integration events. This work provides insight on whether recently reported AAV-derived closed-end DNA and its variants hold promise in achieving non-integrating gene delivery.

## Results

### Design and synthesis of the linear DNA constructs

To investigate the effect of varying the structure of the ends of double stranded linear DNA end region geometry on chromosomal integration, we designed four DNA constructs, each encoding a green fluorescent protein (GFP) reporter and puromycin resistance. We first constructed a circular reporter plasmid and subsequently synthesized plain linear, CELiD and blocked-end constructs using this plasmid as a common backbone, thus minimizing any possible effect originating from differences in sequence or synthesis method. The backbone plasmid consisted of two constitutive expression cassettes for the GFP reporter and puromycin resistant selection marker, in addition to the WPRE element for increasing mRNA stability, all flanked by *BsaI* restriction sites designed to give different four-base overhangs at the two ends (Figure 1A). Plain linear DNA with unmodified ends was created simply by digestion with *BsaI* followed by purification (Figure 1B). Because *BsaI* is a type IIS enzyme that cleaves at a short distance outside of its recognition site, the two sticky ends could have different and non-complimentary sequences, avoiding potential self-ligation. CELiDs were synthesized by further modifying each end of this plain linear DNA by ligation with hairpin-forming oligonucleotides using a Golden Gate assembly (Figure 1B, 1C).

**Figure 1.**
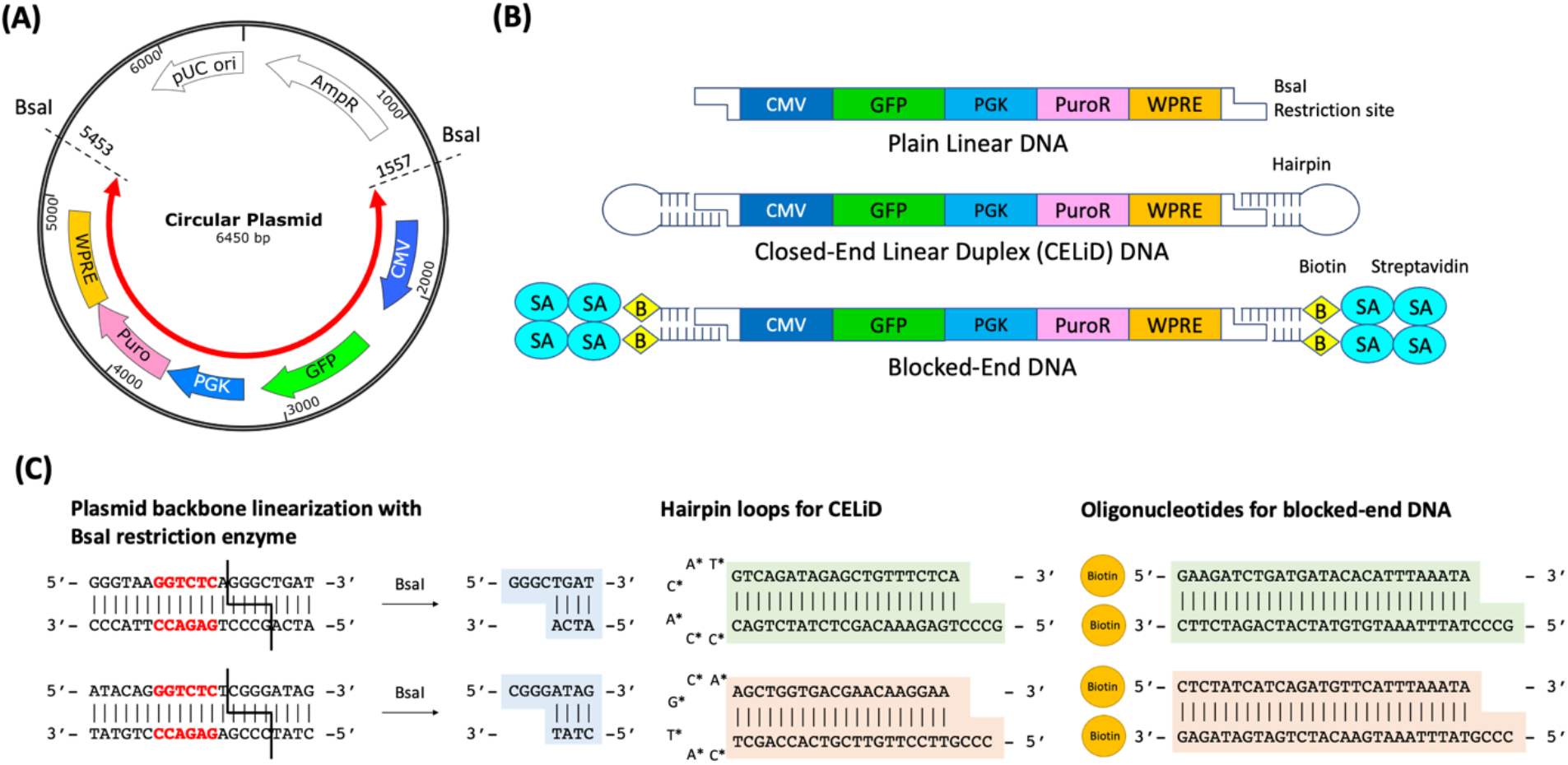
Design of DNA constructs used in this study. **(A)** Circular plasmid used as the common backbone for constructing various end-modified linear DNAs. The plasmid consisted of two constitutive expression cassettes for the GFP reporter and puromycin resistant selection marker, in addition to the WPRE element, flanked by the two *BsaI* restriction sites. The red arrow indicates a portion of the plasmid corresponding to the linear DNAs. **(B)** Structure of the linear DNAs. The end regions of the CELiD consisted of closed hairpin loop structures. The ends of blocked-end DNA contained biotin-labeled oligonucleotides, which were further non-covalently bound to streptavidin tetramers. **(C)** Detailed sequences of the sticky ends created by the plasmid backbone linearization by the *BsaI* restriction enzyme, as well as the hairpin loops and oligonucleotides complementary to each end.

To construct blocked-end DNAs, we first synthesized biotin end-labeled DNAs using the same *BsaI*-digested linear DNA as for the CELiDs, using two different biotin-labeled duplex oligonucleotide fragments containing specific sticky ends corresponding to each end of the linear DNA (Figure 1C, 2A). Two different versions were synthesized. In one version biotin was incorporated only at the 5’ ends (“single biotin”). In the other version, biotin was incorporated in both the 3’ and 5’ ends (“double biotin”). We then tested three types of biotin-binding proteins, avidin, streptavidin, and neutravidin; protein-to-DNA molar ratios were varied from 0.5 to 2 in order to determine the optimal reaction condition. Binding of these proteins to the linear DNA was demonstrated in a gel-shift assay, in which DNA migration in an agarose gel is retarded upon protein conjugation (Figure 2B). Streptavidin showed the strongest binding capacity, with all of the DNA shifting upon addition of equimolar or 2x excess streptavidin. In addition, a band running at a much higher apparent molecular weight was observed, which was likely a circular form in which the streptavidin tetramer is bound to both ends of the DNA. Avidin and neutravidin did not bind as strongly, as inferred from the gel shift assay. Thus, we chose to use streptavidin for constructing blocked-end linear DNA.

**Figure 2.**
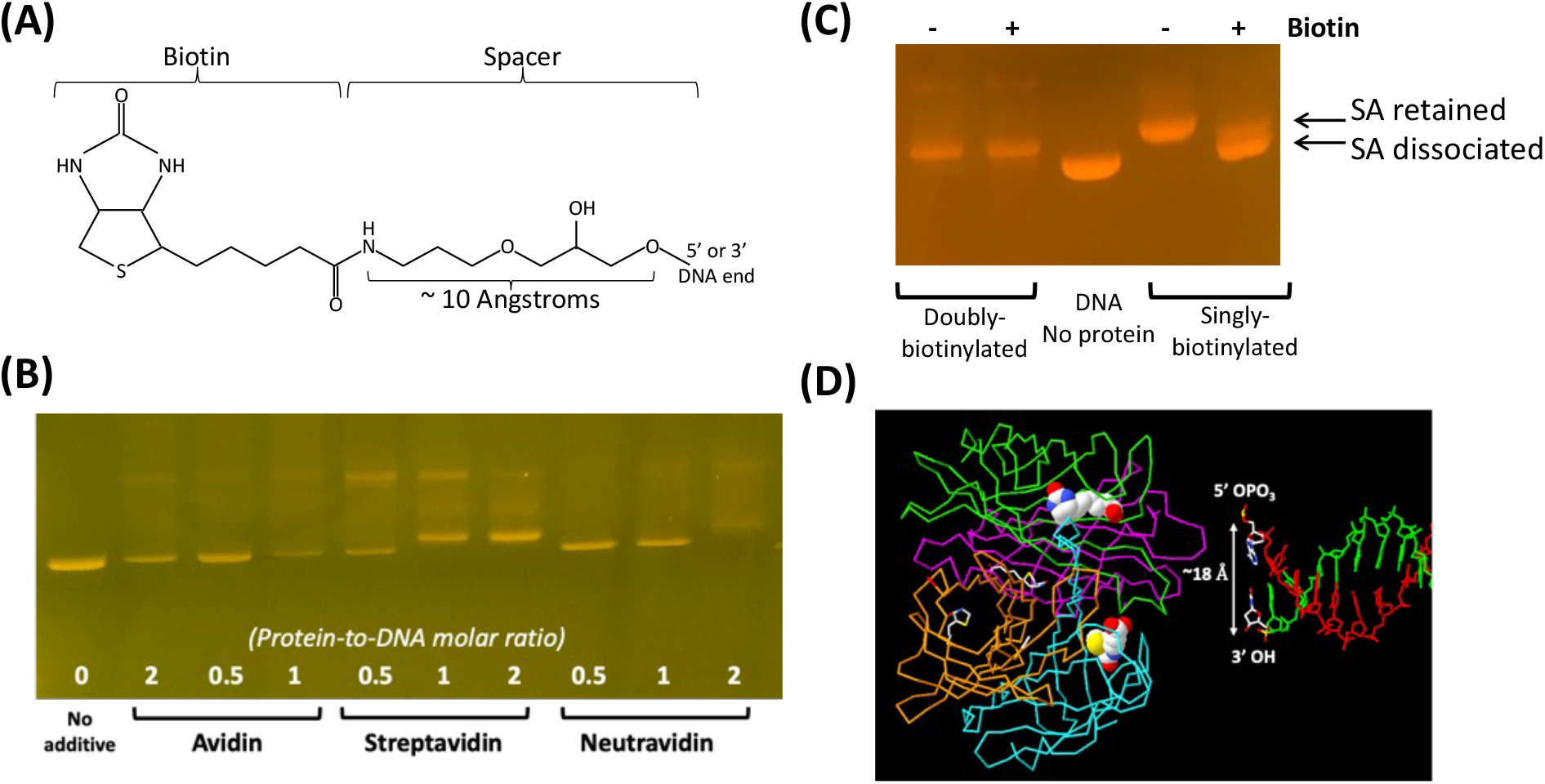
Synthesis and characterization of blocked-end linear DNA. **(A)** Structure of the 5’ end of biotin-labeled linear DNA. [30] **(B)** Agarose gel electrophoresis showing the formation of DNA-protein complexes using three different biotin-binding proteins: avidin, streptavidin and neutravidin. Bound complexes migrate slower compared to unbound DNA. The numbers indicate protein-to-DNA molar ratio, which varied from 0.5 to 2 for each protein. In the streptavidin-bound complexes, the DNA species that migrates slightly slower than the linear DNA alone (leftmost lane) presumably has a streptavidin tetramer bound to each end, while the most slowly migrating species may be a relaxed circle that is held together by a single streptavidin tetramer. **(C)** An agarose gel showing the dissociation of streptavidin (SA) from blocked-end DNA complexes where the DNA has either a biotin at only the 5’ ends or biotins at both the 5’ and 3’ ends, after overnight incubation with (+) and without (-) excess free biotin. Arrows indicate bands corresponding a species with a streptavidin bound to one of the two ends (“SA retained”) and a completely dissociated species, respectively. **(D)** Structural illustration of the end of a ‘blocked-end linear’ DNA, showing a juxtaposition of the end of a DNA fragment with the tetrameric biotin-streptavidin complex (PDB file 1STP). [17] The 5’ and 3’ ends of the DNA backbone are about 18 Angstroms apart, and the distance between the carboxyl groups in the biotins are also about 18 Angstroms.

Next, we tested the stability of our blocked-end constructs by measuring streptavidin dissociation over time under excess free biotin. We assumed that any streptavidin dissociated from the DNA complex would be bound by free biotin, which would then prevent re-binding to the DNA complex. Streptavidin/biotin-DNA complexes using DNAs that were double-biotinylated (5’ and 3’ ends) or single-biotinylated (5’ end only) were incubated with a 10 x molar excess of free biotin at 37°C for various times. Agarose gel electrophoresis showed that after 18 hours of incubation with excess free biotin, the majority of streptavidin was dissociated from the single-biotinylated DNA, whereas the double-biotinylated construct did not show any noticeable detachment (Figure 2C). Both complexes remained tightly bound in the absence of free biotin. This observation suggested that double-biotinylated linear DNA forms a significantly tighter complex with streptavidin than its single-biotinylated counterpart. This may be attributed to the favorable geometry of the double-biotinylated DNA complex, in which the distance between 5’ and 3’ ends (∼18 Angstroms) is approximately similar to the distance between the carboxyl groups of adjacent biotins bound to the streptavidin tetramer according to the known crystal structure [17] (Figure 2D). Hence, double-biotinylated DNA was used to construct blocked-end construct for the rest of this study.

### Comparing integration of DNA constructs upon transfection into human cell line

Having confirmed the formation as well as optimal design of the streptavidin-capped blocked-end linear DNA, we then transfected it into a human cell line and compared the frequency of integration to those of circular, plain linear and CELiD-type DNAs. We transfected the same amount of each DNA construct into human embryonic kidney (HEK) 293 cells using lipofectamine and monitored the changes in percentage of reporter-expressing cells over 3 weeks using flow cytometry (Figure 3A). Cells were split 1:10 every three days, which essentially maintained the cells at a constant density because the doubling time of HEK293 cells is slightly less than 1 day. As the transfected cells were maintained without an antibiotic selection, transfection percentage was expected to decrease (Figure 3B). Cell division would initially dilute the reporter gene concentration, and subsequently produce new cells that do not contain the reporter gene (Figure 3C). Thus, after an extended period of cell growth, transiently transfected reporter expression would become negligible, and those still displaying a fluorescence signal would represent the cell population that had incorporated a chromosomal insertion of the reporter gene.

**Figure 3.**
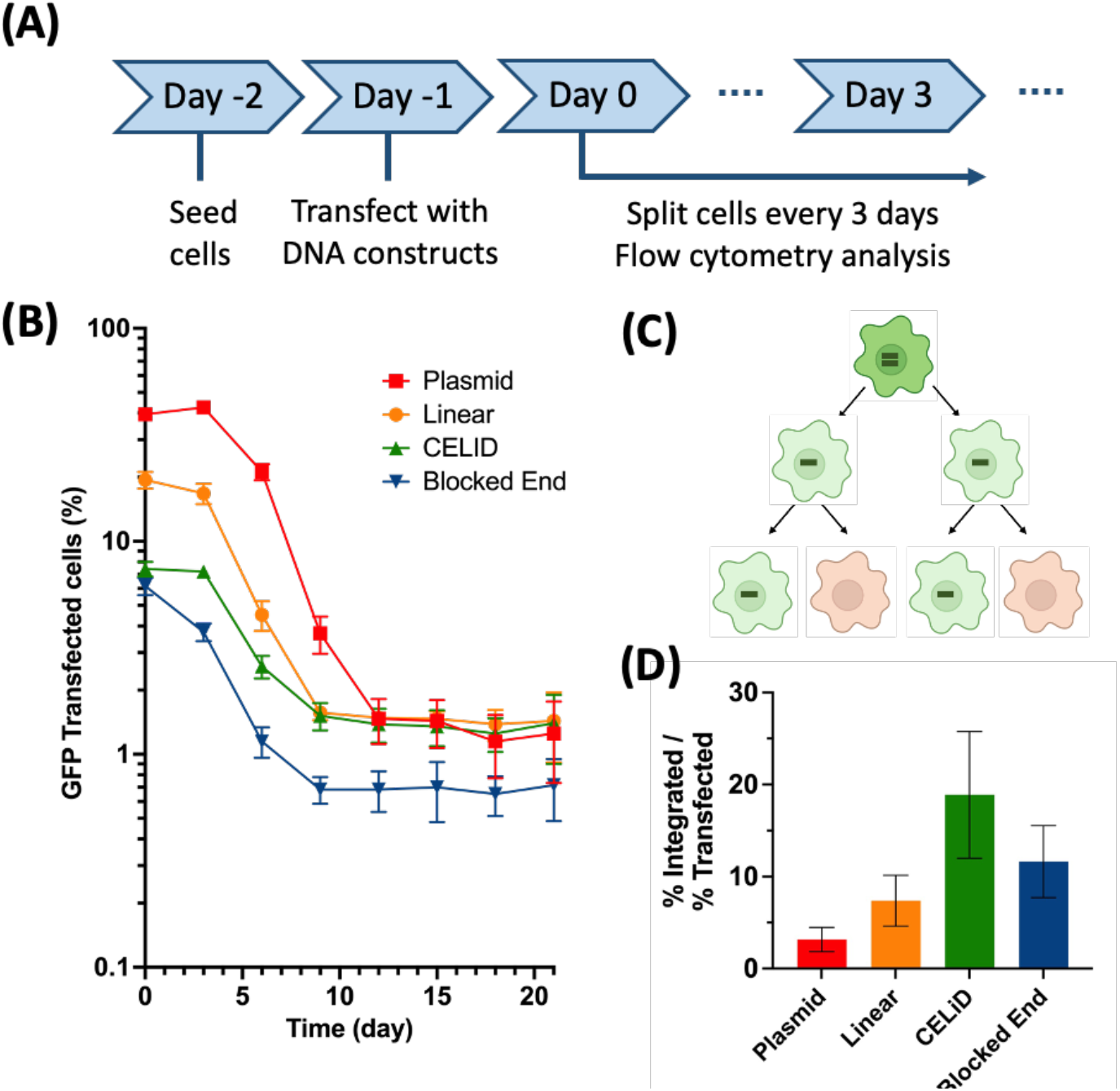
Comparing chromosomal integration rate of various DNA structures upon transfection into HEK293 cells. **(A)** Steps in the experiment. **(B)** Decrease in percentage of GFP expressing cells over time. Cells were split and analyzed by flow cytometry every 3 days until day 21. **(C)** Cartoon showing the segregation of transiently transfected DNAs during cell growth in an absence of drug selection. (D) Normalized integration frequency represented by (% GFP positive cells at day 21) / (% GFP positive cells at day 0).

As expected, cells transfected with all four DNA constructs initially showed a decreasing GFP positive population and subsequently reached a plateau, which represented the percentage of the cells having the reporter gene integrated into their chromosome (Figure 3B). Overall, lipofectamine-mediated transfection with plasmids and various linear DNAs resulted in 6% to 40% of the cells initially expressing GFP. Circular, supercoiled plasmid DNA gave the highest transient transfection frequency, and the relative frequencies were: plasmid > non-modified linear > CELiD > blocked end (Figure S1). After the first split, the proportion of cells expressing GFP was about the same, but the strength of GFP expression was reduced (Figure S2), consistent with the idea that the cells were transfected with multiple DNA molecules that then segregated from each other during cell division. Upon the 2nd and 3rd splits, the proportion of cells expressing GFP was reduced 5-10 fold per split, consistent with the idea that the reporter DNAs are being segregated without replicating, and cells at this stage contain one such DNA. On about day 9-12, the fraction of cells expressing GFP stabilized, formally indicating that the GFP DNAs are replicating as well as not conferring a selective disadvantage to the host cells, and suggesting that these DNAs have integrated. An alternative hypothesis is that a portion of the GFP expression vectors assume an extrachromosomal but replication-competent form; this possibility is addressed below.

Unexpectedly, our observation suggested that the end modification of linear DNA did not significantly decrease the rate of integration. At 24 days after transfection, circular plasmid, non-modified linear DNA and CELiD all showed integration between 1.2 – 1.5% of the total cell population, whereas the blocked end construct showed slightly decreased integration of 0.6%. When we normalize the final percentage of GFP-positive cells at day 24 to the initially transfected percentage at day 0, apparent integration potency was: CELiD > blocked end ∼ non-modified linear > circular plasmid (Figure 3D). Specifically, integration rate of the blocked end construct did not show a statistically significant difference compared to that of non-modified linear DNA (both ∼10%), and the CELiDs even showed an increased tendency for integration (∼20%). Additionally, we observed a consistent result from a biological repeat experiment; despite relatively lower initial transfection efficiencies, the time-dependent decrease in GFP-positive cells and normalized integration frequency for each construct showed a similar trend to support our observation (Figure S3).

Data in Figure 3 suggest that, on average, about 100 copies of transcriptionally active DNA enter a transfected cell. For the plasmid-transfected cells, the proportion of GFP+ cells is relatively steady until after the second 1:10 split, suggesting that 100 plasmid DNA molecules present in an initially transfected cell are distributed into 100 cells at this point. According to this idea, the initially transfected cells go through six to seven cell divisions, during which non-replicating DNA is passively segregated into daughter cells. These results further suggest that in the plasmid-transfected cells, the fraction of plasmids that integrate is about 10^−4^ of the input plasmids. The quantitation of GFP expression in the transfected cells (Figure S2) does not indicate that silencing of transgene gene expression occurs.

To verify that the GFP-positive cell population after reaching plateau actually contains the reporter sequence in genomic DNA, we isolated the clones of individual fluorescent cells and analyzed their genomes for the presence of the GFP gene using qPCR-based copy number assay. For each type of transfected DNA, two separate monoclonal cells were isolated from the population of cells stably expressing GFP (Figure 4A). qPCR reactions specific to the GFP reporter revealed that all clones contained the target sequence, and that their copy numbers varied from 1 to 3 copies (Figure 4B). In addition, inverse PCR experiment identified at least one integration junction from each monoclonal cell genome to support our result (Figure 4C). Thus, we confirmed that the lipofectamine-mediated transfection with plasmids and various linear DNAs had indeed led to the stable incorporation of genes into the chromosomes.

**Figure 4.**
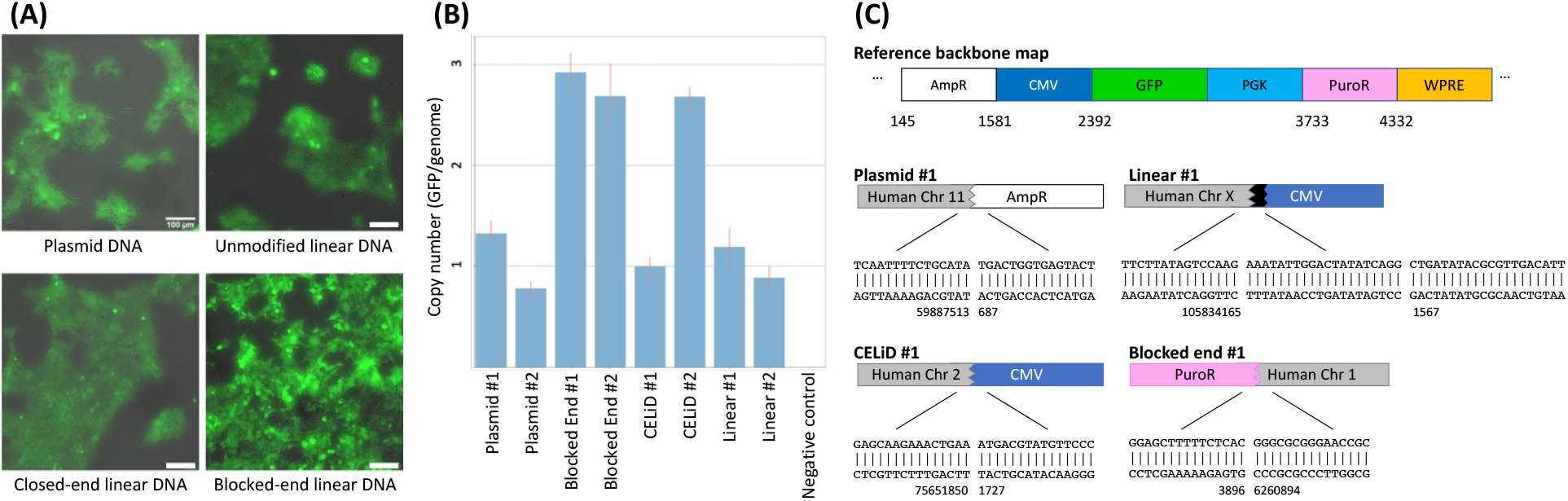
Gene integration analysis using cloned cells. GFP+ cells from the experiment in Figure 3B at day >20 were isolated by FACS and plated at limiting dilution to obtain clones. **(A)** Images of monoclonal cells isolated from HEK293 cells transfected with each type of DNA construct. Images taken from brightfield and GFP channel were overlapped. Scale bars indicate 100 µm. **(B)** Copy number analysis of monoclonal cells transfected with different DNA types. For each DNA type, genomic DNA from two separate clonal cell lines was extracted and analyzed by qPCR using probes specific to the GFP gene. **(C)** Detailed sequences of the integration junctions identified for each type of monoclonal cells using an inverse PCR method.

## Discussion

The frequency at which transgenes spontaneously integrate into a mammalian chromosome is centrally important in understanding the safety of gene therapy systems. Retroviruses were the first gene therapy delivery system, but the high integration frequency and ability to transactivate nearby genes, including oncogenes, has led to leukemia in patients. [3] Subsequent retroviral vectors have been engineered to eliminate transactivation, but this still leaves the potential problem of insertion into tumor suppressors. An alternative strategy has been to use “non-integrating” vectors such as Adeno-Associated Virus (AAV), or Closed-End Linear DNAs. Unlike retroviruses, AAV DNAs are not obligated to integrate during the virus life cycle, [7] so this virus is dubbed “non-integrating.” However, the dose of AAV used in gene therapy clinical trials is quite high, so even a low rate of integration could pose a risk of insertion into tumor suppressor genes. Moreover, in a mouse model, administration of AAV leads to oncogenic integration events. [8, 9]

We quantitated the frequency of spontaneous unselected chromosomal integration of transgenes that were delivered into cultured mammalian cells as covalently closed plasmids or in three different linear formats: DNA with exposed 5’ and 3’ ends, Closed-End Linear DNAs (CELiDs), and DNA with biotinylated 5’ and 3’ ends further capped with streptavidin (Figure 2). We developed the streptavidin-capped DNAs to try to minimize the potential for non-homologous end joining (NHEJ). Within streptavidin, the biotin binding sites are the same distance apart as the 3’ and 5’ ends of DNA (∼18 Angstroms), and dissociation of streptavidin from doubly biotin capped DNA was undetectable (Figure 2C).

Several lines of evidence indicate that spontaneous, unselected integration occurred at a high rate. First, after transfection, the fraction of cells expressing GFP stabilized at 1-20% of the initially transfected cells; this fraction was stable over at least three to four 10-fold splits. This is consistent with either integration or conversion into a replication-competent form, such as a long concatemer. Second, when clones of GFP+ cells are isolated after FACS and extreme limiting dilution, quantitative PCR indicated that only one to three copies of transgene DNA are present. Finally, characterization of several cell clones identified junctions between transgene DNA and human chromosomal DNA.

Different forms of DNA integrate at different frequencies. After transfection into HEK293 cells, about 1-10% of the transiently transfected cells became stably transfected (Figure The circular plasmid DNA has a lower rate of integration than all of the linear forms, which show comparable integration frequencies regardless of the configuration of the DNA ends. We also estimated that transfected cells received about 100 functional expression units per cell, based on the fact that the proportion of GFP-expressing cells stays roughly constant through two 10-fold splits before decreasing. Thus, the integration frequency is about 10^−4^/DNA for circular plasmids and about 10^−3^ for linear DNAs. These results are consistent with a previous report, [11] in which mice *in vivo*-transfected with CELiDs by hydrodynamic injection demonstrated more stable expression of the transgene in the liver, compared to expression in livers *in vivo*-transfected with a circular plasmid.

The linear DNAs all integrated at about the same frequency, and when we could identify junctions between human and plasmid DNA, the ends of the plasmid DNA were quite far from the end of the transfected DNA where it was cut by the restriction enzyme for linearization (Figure 4C). The integration of these diverse forms of linear DNA suggests that the cell has a mechanism for eliminating non-natural DNA ends and then rescuing the DNA by integration.

These results may have implications that for AAV gene therapy, wherein viral particles deliver a mostly single-stranded DNA with closed hairpin ends. Upon delivery to the nucleus, such DNAs are converted into a CELiD form. [18, 19] Clinical trials of AAV-mediated delivery of genes for hemophilia A and B (Factor VIII and Factor IX deficiency, respectively) illustrate that the doses are high enough to result in integration events in almost every cell, assuming efficient infection. George *et al*. used doses of 5×10^11^ to 1.5×10^12^ vector genomes per kg body weight in a Phase I-II study of Factor VIII delivery by AAV. [20, 21] These correspond to absolute doses of 3.5×10^13^ to 10^14^ vector genomes for a 70-kg person. The adult human liver contains about 1.5×10^11^ cells (10^8^ cells/gm; Liver mass is ∼1.5 kg). [22, 23] Thus, the ratio of AAV transducing genomes to liver cells is about 100-1,000:1, and if the integration rate is the same as what we observed for CELiDs in this work, the fraction of liver cells experiencing an integration event would be 10-100%, assuming every AAV particle infects a cell. One important difference between our experiments and human AAV gene therapy is that HEK293 cells are rapidly dividing and liver cells are not, which may lead to differences in integration rates.

Our analysis is consistent with observations from studies reporting transgene integration and tumorigenesis in animals injected with AAV vectors. [7-9, 24-28] In comparing the safety of retroviral and AAV-based vector systems, these calculations suggest that the high doses used in AAV therapies may nullify the advantage deriving from the fact that AAV is a “non-integrating” vector.

Our results suggest that AAV-based gene therapy may be more susceptible to integration and subsequent insertional mutagenesis than previously thought. The results also indicated that engineering end regions of linear DNA is insufficient to prevent its insertion into host cell genome. Recent findings from long-term study of AAV gene therapy consistently indicate evidence of genomic integration. Currently, gene therapy aims to treat cancer or rare genetic diseases where long-term risks are acceptable. Further development of non-integrating gene delivery methods may need to precede wider application of gene therapy.

## Methods

### Cell culture and maintenance

Dulbecco’s Modified Eagle’s Medium (DMEM) (ATCC 30-2002), Heat-inactivated Fetal Bovine Serum (FBS) (Thermo Fisher Scientific 10082147), and Penicillin-Streptomycin (Corning 30-002-Cl) were all vacuum sterilized by fi0ltration (0.22 μm pore size, Corning, 430767) and used for maintaining cell growth. Homo sapiens HEK-293T cells (ATCC, CRL-3216) were maintained in DMEM supplemented by FBS and Penicillin-Streptomycin at 37°C. Once the cells reached 70% confluency, they were split into new flasks at one tenth of the density using trypsin (Trypsin EDTA 1X 0.25% Trypsin/ 2.21mM EDTA in HBSS without sodium bicarbonate, calcium and magnesium, VWR 45000-664).

### Circular and Linear DNA synthesis

Original circular plasmid DNA containing GFP reporter under CMV promoter, Puromycin resistance cassette under PGK promoter and WPRE sequence was ordered from Twist Biosciences. Unmodified plain linear DNA was synthesized by first digesting this plasmid with a *BsaI* restriction enzyme (New England Biolab), followed by separation in agarose gel and purification using the gel extraction kit (Qiagen).

### CELiD DNA synthesis

Two hairpin oligos were ordered from Integrated DNA Technologies (IDT). Their sequences were:

5’-GCCCTGAGAAACAGCTCTATCTGAC*C*A*C*A*T*GTCAGATAGAGCTGTTTCTCA-3’

5’-CCCGTTCCTTGTTCGTCACCAGCTC*A*T*G*C*A*GAGCTGGTGACGAACAAGGAA-3’ where the asterisks indicate thioester linkages in the loop sections. Plain linear DNA was incubated with 10x molar excess of each hairpin oligo in a Golden Gate Assembly (NEB). Subsequently, excess hairpin oligos were digested using Exo III (NEB) and synthesized CELiDs were purified using a PCR cleanup kit (Qiagen).

### Blocked-end DNA synthesis

Four Biotinylated oligonucleotides forming two double-stranded biotinylated fragments were ordered from IDT. Their sequences were:

5’Bio-CTCTATCATCAGATGTTCATTTAAATA-3’

5’-CCCGTATTTAAATGAACATCTGATGATAGAG-Bio3’

5’Bio-GAAGATCTGATGATACACATTTAAATA-3’

5’-GCCCTATTTAAATCTGTATCATCAGATCTTC-Bio3’

Each pair was mixed, heated at 95°C and cooled slowly to anneal the two strands. Similar to the CELiD synthesis, plain linear DNA was incubated with 10x molar excess of each biotinylated fragment in a Golden Gate Assembly. Resulting biotinylated DNA was then further incubated with 2x molar excess of streptavidin for 10 min, followed by Exo III treatment to digest excess oligos and purification.

### Cell seeding

Prior to seeding, 96-well cell culture plate was coated with Purecol collagen solution (Advanced Biomatrix) diluted 1:30 with Phosphate-Buffered Saline (PBS) (Thermo Fisher Scientific, AM9625). Each well was incubated with 100 µL collagen for 1 hr and rinsed with 150 µL PBS. HEK-293T cells were washed with Phosphate-Buffered Saline (PBS) (Thermo Fisher Scientific, AM9625) and trypsinized (Trypsin EDTA 1X 0.25% Trypsin/ 2.21mM EDTA in HBSS without sodium bicarbonate, calcium and magnesium, VWR 45000-664). Cell density was determined by optical density with an automated cell counter (Bio-Rad TC20), cell counting slides (Bio-Rad, 1450015) and 0.4% solution Trypan Blue (Bio-Rad, 1450021). After live cell count was obtained, cells were diluted to 2.7 × 10^5^ cells per mL in DMEM supplemented with 10% FBS, and 150 μL of cells were added to each well to seed 4 × 10^4^ cells per well. Plates were then incubated at 37°C for 24 hours.

### Transfection of cells with various DNA constructs

HEK-293T cells were transfected 24 hours after plating on 96-well plates and were carried out in DMEM supplemented with 10% FBS. Per well, 50 ng of DNA construct was transfected using lipofectamine 3000 kit (Thermo Fisher Scientific, L3000015). Each well was transfected with 10 μL of the Opti-MEM (Thermo 31985062) media mixture containing 50 ng of DNA, 0.15 μL lipofectamine reagent, 0.2 μL p3000 reagent. For each construct, total of six wells were transfected as replicates. Media was exchanged to fresh DMEM supplemented with 10% FBS 24 hrs after the addition of transfection reagent. Subsequently, 24 hours later, cells were split 1:10 using trypsin-EDTA into a new 96-well plate, and remaining cells were used for the flow cytometry analysis. Similarly, cells were split 1:10 every three days during the 24-day experiment period.

### Flow cytometry

Cells in a 96-well plate were trypsinized and resuspended in flow cytometry buffer (Invitrogen). Fluorescence was measured on a LSR II flow cytometer equipped with a HTS sampler (BD Biosciences) using the following filter configuration: excitation, 488 nm; emission, 530/30.

### Microscopy imaging

Images were taken using a Nikon Eclipse Ti-2 widefield inverted microscope equipped with Hamamatsu Flash 4.0 LT camera at 20x objective. For imaging green fluorescence, the following filter configuration was used: excitation, 466/40; emission, 525/50.

### Isolation of monoclonal cell population

To generate individual clones of the fluorescent cells, transfected cells at day 24 were first sorted using a FACSAria II cell sorter (BD Biosciences) using the same filter configuration as above. Then, monoclonal cells were isolated from the sorted GFP-positive population using limiting dilution method. In specific, homogenized cells were diluted up to concentration of 2.5 cells/mL, and 100 μL of this cell solution was transferred into each well of a 96-well plate. At this ratio, one out of four wells contained a single cell that should grow into monoclonal population. Seeded cells were cultured for 14 days, checked for fluorescence, and expanded into larger culture dishes. On average, 5-10 clones were successfully isolated, which were less than expected; such result may be attributed to the low cell viability at a very low density.

### Copy number assay and inverse PCR

Genomic DNA was extracted from monoclonal cells using DNeasy Blood and Tissue kit (Qiagen) following the instructions. Subsequently, qPCR-based copy number assay was carried out using Taqman Copy Number Assay and RNAse P Reference Assay (Thermo Fisher Scientific). qPCR probes specific to the GFP reporter gene was custom ordered from Thermo Fisher Scientific. Reactions were run using Quantstudio 7 Real-Time PCR system (Applied Biosystems), and subsequently analyzed for copy number using CopyCaller software (Applied Biosystems). For the inverse PCR, extracted genomic DNA was digested using a range of restriction enzymes including *EcoRI, BamHI, SphI* and *NcoI* (NEB) and ligated using T4 DNA ligase (NEB). Then, ligated circular DNAs were amplified using GFP reporter-specific primers (IDT) and Q5 hot start DNA polymerase (NEB). Amplified DNA was sequenced (Azenta) to identify the integration junction. Acquired sequences were compared with human genome sequence provided by UCSC Genome Browser (http://genome.uscs.edu), [29] using a genome assembly released in Dec 2013.

## Acknowledgements

This research was funded by the DARPA (DARPA-STTR Phase I Contract 140D0420C0057 and DARPA-STTR Phase II Contract HR001121C0142)

## Author contributions

S. L. designed the research, performed the experiments, analyzed the data and wrote the paper. R. Y. designed, synthesized and characterized the DNA constructs. J.W. and P.S. analyzed the data, contributed to experiment design and edited the paper.

## Data availability statement

The datasets generated during and/or analyzed during the current study are either included in this published article (supporting information), or available from the corresponding author on reasonable request.

## Additional information

J. W. and R. Y are employed by General Biologics, Inc., a for-profit corporation, and P. S. is a founder of General Biologics, Inc. S. L. declares no competing financial interests.

