## Supporting Information for "High spontaneous integration rates of end-modified linear DNAs upon mammalian cell transfection"

Samuel Lim*[1,2], R. Rogers Yocum[3], Pamela A Silver[1,2], Jeffrey C Way*[3]

**Author affiliations**

1. Department of Systems Biology, Harvard Medical School, Boston, Massachusetts 02115, United States.
2. Wyss Institute for Biologically Inspired Engineering, Boston, Massachusetts 02115, United States.
3. General Biologics, Inc., 108 Fayerweather Street, Unit 2, Cambridge, Massachusetts 02138, United States.

**Author contacts**

Samuel Lim:

R. Rogers Yocum:

Jeffrey C Way:

Pamela A Silver:

*Corresponding authors

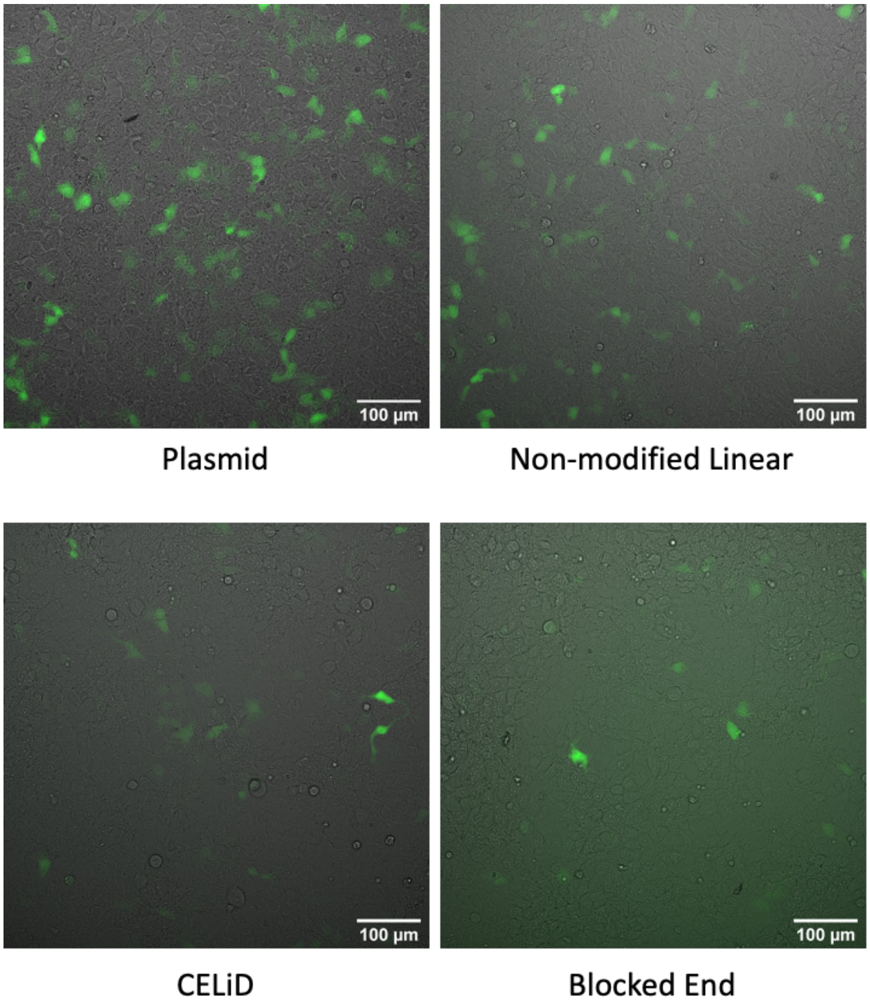


**Figure S1.** Images of the HEK293 cells transfected with each type of DNA construct 24 hours after transfection. Images taken from brightfield and green fluorescent channel were overlapped. Scale bar indicates 100 µm.

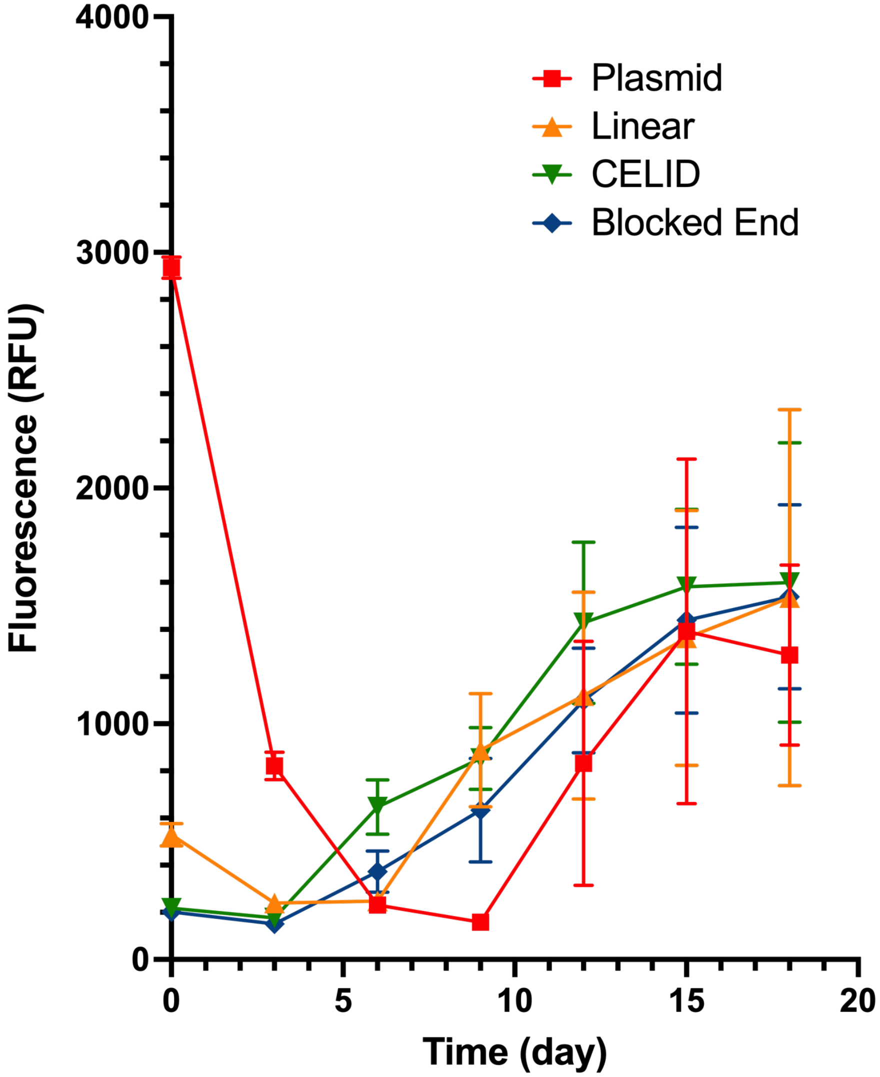


**Figure S2.** Change in fluorescence intensity of transfected cells over time. Median fluorescence intensity of the GFP positive cell population was measured from day 0 to 21, using flow cytometry. In the cells transfected with plasmid and linear DNAs, the cellular expression levels in GFP+ cells increases significantly when these cells become a stabilized fraction of the total. This result suggests that the integration events occur in chromosomal loci that promote particularly strong expression.

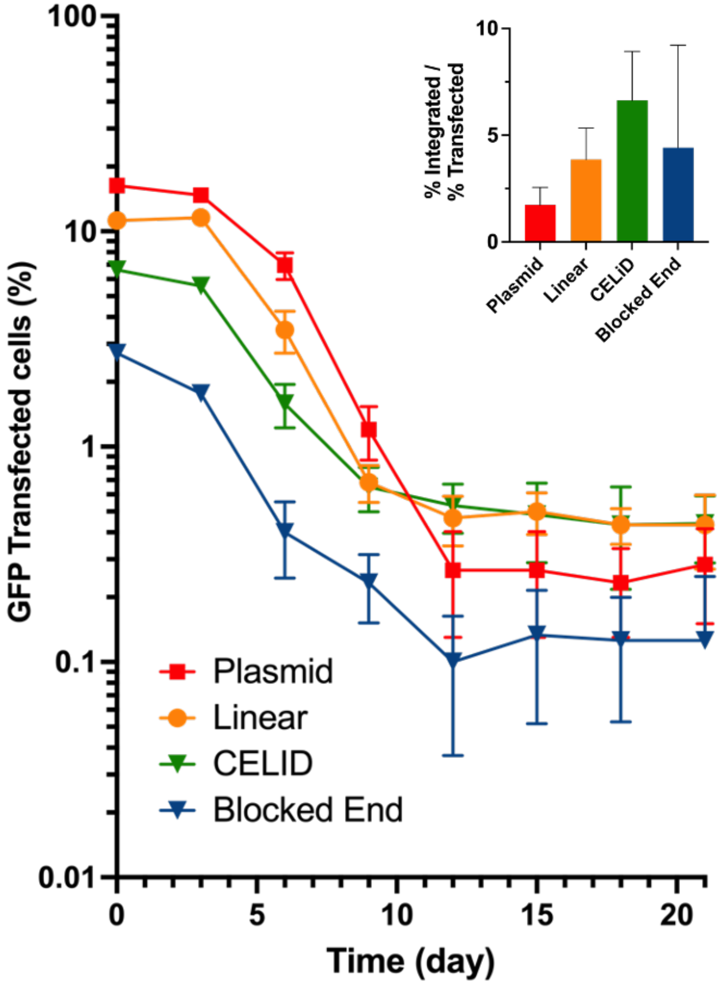


**Figure S3.** A biological repeat of the DNA transfection experiment. Results were consistent with those described in Figure 3.

```
LOCUS       Exported                6450 bp DNA     circular SYN 12-DEC-2022
DEFINITION  synthetic circular DNA
ACCESSION   .
VERSION     .
KEYWORDS    .
SOURCE      synthetic DNA construct
  ORGANISM  synthetic DNA construct
REFERENCE   1  (bases 1 to 6450)
  AUTHORS   .
  TITLE     Direct Submission
  JOURNAL   Exported Dec 17, 2022 from SnapGene 6.0.5
            https://www.snapgene.com
FEATURES             Location/Qualifiers
     source          1..6450
                     /note="/dnas_title=pRG49 figure"
                     /organism="synthetic DNA construct"
     misc_feature    complement(145..1005)
                     /locus_tag="AmpR"
                     /label=AmpR
     misc_feature    1557..1562
                     /note="BsaI recog"
     misc_feature    1581..2164
                     /label=CMV promoter
                     /note="CMV promoter"
     misc_feature    2392..3111
                     /locus_tag="GFP"
                     /label=GFP
     misc_feature    3221..3720
                     /locus_tag="PGK prom."
                     /label=mPGK
                     /label=PGK prom.
     misc_feature    3733..4332
                     /note="purR codon optimized"
     misc_feature    4334..4930
                     /locus_tag="WPRE"
                     /label=WPRE
     misc_feature    5000..5234
                     /locus_tag="3' dLTR"
                     /label=3' dLTR
     misc_feature    5306..5437
                     /locus_tag="SV40 PolyA"
                     /label=SV40 PolyA
                     /note="/dnas_title=Ter"
     misc_feature    5453..5458
                     /note="BsaI recog"
     misc_feature    complement(5847..6450)
                     /locus_tag="pUC ori"
                     /label=pUC ori
ORIGIN
        1 gtctgacgct cagtggaacg aaaactcacg ttaagggatt ttggtcatga gattatcaaa
       61 aaggatcttc acctagatcc ttttaaatta aaaatgaagt tttaaatcaa tctaaagtat
      121 atatgagtaa acttggtctg acagttacca atgcttaatc agtgaggcac ctatctcagc
      181 gatctgtcta tttcgttcat ccatagttgc ctgactcccc gtcgtgtaga taactacgat
      241 acgggagggc ttaccatctg gccccagtgc tgcaatgata ccgcgagacc cacgctcacc
      301 ggctccagat ttatcagcaa taaaccagcc agccggaagg gccgagcgca gaagtggtcc
      361 tgcaacttta tccgcctcca tccagtctat taattgttgc cgggaagcta gagtaagtag
      421 ttcgccagtt aatagtttgc gcaacgttgt tgccattgct acaggcatcg tggtgtcacg
      481 ctcgtcgttt ggtatggctt cattcagctc cggttcccaa cgatcaaggc gagttacatg
      541 atcccccatg ttgtgcaaaa aagcggttag ctccttcggt cctccgatcg ttgtcagaag
      601 taagttggcc gcagtgttat cactcatggt tatggcagca ctgcataatt ctcttactgt
      661 catgccatcc gtaagatgct tttctgtgac tggtgagtac tcaaccaagt cattctgaga
      721 atagtgtatg cggcgaccga gttgctcttg cccggcgtca atacgggata ataccgcgcc
      781 acatagcaga actttaaaag tgctcatcat tggaaaacgt tcttcggggc gaaaactctc
```

841 aaggatctta ccgctgttga gatccagttc gatgtaaccc actcgtgcac ccaactgatc
901 ttcagcatct tttactttca ccagcgtttc tgggtgagca aaaacaggaa ggcaaaatgc
961 cgcaaaaaag ggaataaggg cgacacggaa atgttgaata ctcatactct tcctttttca
1021 atattattga agcatttatc agggttattg tctcatgagc ggatacatat ttgaatgtat
1081 ttagaaaaat aaacaaatag gggttccgcg cacatttccc cgaaaagtgc cacctgacgt
1141 ctaagaaacc attattatca tgacattaac ctataaaaat aggcgtatca cgaggccctt
1201 tcgtctcgcg cgtttcggtg atgacggtga aaacctctga cacatgcagc tcccggagac
1261 ggtcacagct tgtctgtaag cggatgccgg gagcagacaa gcccgtcagg gcgcgtcagc
1321 gggtgttggc gggtgtcggg gctggcttaa ctatgcggca tcagagcaga ttgtactgag
1381 agtgcaccat atgcggtgtg aaataccgca cagatgcgta aggagaaaat accgcatcag
1441 gcgccattcg ccattcaggc tgcgcaactg ttgggaaggg cgatcggtgc gggcctcttc
1501 gctattacgc cagctggcga aagggggatg tgctgcaagg cgattaagtt gggtaaggtc
1561 tcagggctga tatacgcgtt gacattgatt attgactagt tattaatagt aatcaattac
1621 ggggtcatta gttcatagcc catatatgga gttccgcgtt acataactta cggtaaatgg
1681 cccgcctggc tgaccgccca acgacccccg cccattgacg tcaataatga cgtatgttcc
1741 catagtaacg ccaataggga ctttccattg acgtcaatgg gtggactatt tacggtaaac
1801 tgcccacttg gcagtacatc aagtgtatca tatgccaagt acgcccccta ttgacgtcaa
1861 tgacggtaaa tggcccgcct ggcattatgc ccagtacatg accttatggg actttcctac
1921 ttggcagtac atctacgtat tagtcatcgc tattaccatg gtgatgcggt tttggcagta
1981 catcaatggg cgtggatagc ggtttgactc acggggattt ccaagtctcc accccattga
2041 cgtcaatggg agtttgtttt ggcaccaaaa tcaacgggac tttccaaaat gtcgtaacaa
2101 ctccgcccca ttgacgcaaa tgggcggtag gcgtgtacgg tgggaggtct atataagcag
2161 agctctctgg ctaactagag aacccactgc ttagtcgatc ctaccatcca ctcgacacac
2221 ccgccagcgg ccgctcgcca ccatggtaag tgtgaatcga agcgcggctt cagaataccg
2281 ttttggctac aggatacaaa gcccatcgtg gtcctcagat atcgtgcacg tagagttgca
2341 ccgcacgcag tggctaactt ctctctttct ctctccctcc ctgtctttca ggctagcaaa
2401 ggagaagaac tcttcactgg agttgtccca attcttgttg aattagatgg tgatgttaac
2461 ggccacaagt tctctgtcag tggagagggt gaaggtgatg caacatacgg aaaacttacc
2521 ctgaagttca tctgcactac tggcaaactg cctgttccgt ggccgacact agtgacgacg
2581 ctctgctatg gcgtccagtg cttttcaaga tacccggatc acatgaaacg gcatgacttt
2641 ttcaagagtg ccatgcccga aggttatgta caggaaagga ccatcttctt caaagatgac
2701 ggcaactaca agacacgtgc tgaagtcaag tttgaaggtg ataccettgt taatagaatc
2761 gagttaaaag gtattgactt caaggaagat ggcaacattc tgggacacaa attggaatac
2821 aactataact cacacaatgt atacatcatg gcagacaaac aaaagaatgg aatcaaagtg
2881 aacttcaaga cccgccacaa cattgaagat ggaagcgttc aactagcaga ccattatcaa
2941 caaaatactc caattggcga tggccctgtc cttttaccag acaaccatta cctgtccaca
3001 caatctgccc tttcgaaaga tcccaacgaa aagagagatc acatggtcct tcttgagttt
3061 gtaacagctg ctgggattac acatggcatg gatgaactat acaaatcctg agttgtaaat
3121 gagcacacaa aatacacatg ctaaaatatt atattctatg acctttataa aatcaaccaa
3181 aatcttcttt ttaataactt tagtatcaat aattaattaa gggtagggga ggcgcttttc
3241 ccaaggcagt ctggagcatg cgctttagca gccccgctgg gcacttggcg ctacacaagt
3301 ggcctctggc ctcgcacaca ttccacatcc accggtaggc gccaaccggc tccgttcttt
3361 ggtggcccct tcgcgccacc ttctactcct cccctagtca ggaagttccc ccccgccccg
3421 cagctcgcgt cgtgcaggac gtgacaaatg gaagtagcac gtctcactag tctcgtgcag
3481 atggacagca ccgctgagca atggaagcgg gtaggccttt ggggcagcgg ccaatagcag
3541 ctttgctcct tcgctttctg ggctcagagg ctgggaaggg gtgggtccgg gggcgggctc
3601 aggggcgggc tcaggggcgg ggcgggcgcc cgaaggtcct ccggaggccc ggcattctgc
3661 acgcttcaaa agcgcacgtc tgccgcgctg ttctcctctt cctcatctcc gggcctttcg
3721 gatatcgcca ccatgaccga gtacaagccc accgtgcgac tggcaacacg agacgatgta
3781 cccagagcag taagaacgct cgctgctgcg ttcgccgact acccagctac acggcacaca
3841 gttgacccag accgacatat tgaacgggtc accgaactcc aggagctttt tctcactcgc
3901 gtggggctcg atatcggtaa agtttgggtg gccgacgatg gagccgcggt tgcagtgtgg
3961 acaacaccgg aatcagttga agccggtgcg gtttttgcgg aaattggccc acgaatggcg
4021 gaactttccg gatcccgctt ggccgcgcag cagcagatgg agggacttct tgctccgcat
4081 aggccgaaag aaccagcttg gttccttgcc acagttggtg tgtcacctga tcatcaaggc
4141 aaggggctgg ggtctgccgt tgttctgcca ggcgtggagg ccgcggaaag ggctggggtg
4201 cctgccttct tggagacttc agctccacga aatctcccgt tttatgaacg cctgggattt
4261 actgttaccg ctgatgtgga ggttcctgaa ggtcctcgaa cttggtgtat gactaggaaa
4321 ccaggcgcct gagtcgacaa tcaacctctg gattacaaaa tttgtgaaag attgactggt
4381 attcttaact atgttgctcc ttttacgcta tgtggatacg ctgctttaat gcctttgtat
4441 catgctattg cttcccgtat ggctttcatt ttctcctcct tgtataaatc ctggttgctg
4501 tctctttatg aggagttgtg gcccgttgtc aggcaacgtg gcgtggtgtg cactgtgttt
4561 gctgacgcaa cccccactgg ttggggcatt gccaccacct gtcagctcct ttccgggact

```
     4621 ttcgctttcc ccctccctat tgccacggcg gaactcatcg ccgcctgcct tgcccgctgc
     4681 tggacagggg ctcggctgtt gggcactgac aattccgtgg tgttgtcggg gaaatcatcg
     4741 tcctttcctt ggctgctcgc ctgtgttgcc acctggattc tgcgcgggac gtccttctgc
     4801 tacgtccctt cggccctcaa tccagcggac cttccttccc gcggcctgct gccggctctg
     4861 cggcctcttc cgcgtcttcg ccttcgccct cagacgagtc ggatctccct ttgggccgcc
     4921 tccccgcctg gtacctttaa gaccaatgac ttacaaggca gctgtagatc ttagccactt
     4981 tttaaaagaa aaggggggac tggaagggct aattcactcc caacgaagat aagatctgct
     5041 ttttgcttgt actgggactc tctggttaga ccagatctga gcctgggagc tctctggcta
     5101 actagggaac ccactgctta agcctcaata aagcttgcct tgagtgcttc aagtagtgtg
     5161 tgcccgtctg ttgtgtgact ctggtaacta gagatccctc agaccctttt agtcagtgtg
     5221 gaaaatctct agcagtagta gttcatgtca tcttattatt cagtatttat aacttgcaaa
     5281 gaaatgaata tcagagagtg agaggaactt gtttattgca gcttataatg gttacaaata
     5341 aagcaatagc atcacaaatt tcacaaataa agcatttttt tcactgcatt ctagttgtgg
     5401 tttgtccaaa ctcatcaatg tatcttatca tgtctggctc tagctatccc gagagaccct
     5461 gtatccgctc acaattccac acaacatacg agccggaagc ataaagtgta aagcctgggg
     5521 tgcctaatga gtgagctaac tcacattaat tgcgttgcgc tcactgcccg ctttccagtc
     5581 gggaaacctg tcgtgccagc tgcattaatg aatcggccaa cgcgcgggga gaggcggttt
     5641 gcgtattggg cgctcttccg cttcctcgct cactgactcg ctgcgctcgg tcgttcggct
     5701 gcggcgagcg gtatcagctc actcaaaggc ggtaatacgg ttatccacag aatcagggga
     5761 taacgcagga aagaacatgt gagcaaaagg ccagcaaaag gccaggaacc gtaaaaaggc
     5821 cgcgttgctg gcgtttttcc ataggctccg cccccctgac gagcatcaca aaaatcgacg
     5881 ctcaagtcag aggtggcgaa acccgacagg actataaaga taccaggcgt ttccccctgg
     5941 aagctccctc gtgcgctctc ctgttccgac cctgccgctt accggatacc tgtccgcctt
     6001 tctcccttcg ggaagcgtgg cgctttctca tagctcacgc tgtaggtatc tcagttcggt
     6061 gtaggtcgtt cgctccaagc tgggctgtgt gcacgaaccc cccgttcagc ccgaccgctg
     6121 cgccttatcc ggtaactatc gtcttgagtc caacccggta agacacgact tatcgccact
     6181 ggcagcagcc actggtaaca ggattagcag agcgaggtat gtaggcggtg ctacagagtt
     6241 cttgaagtgg tggcctaact acggctacac tagaaggaca gtatttggta tctgcgctct
     6301 gctgaagcca gttaccttcg gaaaaagagt tggtagctct tgatccggca aacaaaccac
     6361 cgctggtagc ggtggttttt ttgtttgcaa gcagcagatt acgcgcagaa aaaaaggatc
     6421 tcaagaagat cctttgatct tttctacggg
//
```

**Figure S4.** The sequence of the plasmid in Figure 1A, represented in GenBank format.
